# Post-encoding slow-wave amplitude during a daytime nap predicts pattern completion from sparse visual cues

**DOI:** 10.64898/2026.07.17.738149

**Authors:** A. Hanert, Y. Fiedler, J. Hacker, P. Vieweg, A. Pedersen, J. Born, A. Burgalossi, T. Bartsch

## Abstract

Pattern completion refers to the reinstatement of a stored memory representation from partial or degraded cues. In this sense, it enables a form of cue-based generalization: the same memory representation can be retrieved across different, incomplete versions of the original input. Sleep supports hippocampus-dependent memory consolidation and may facilitate such flexible retrieval, but it remains unclear whether post-encoding sleep improves visual pattern completion from degraded cues. Previous sleep studies have mainly examined mnemonic discrimination or relational memory, leaving open whether sleep directly enhances the recovery of learned visual scenes from sparse perceptual information.

We tested this question using the Memory Image Completion (MIC) task in a polysomnographic within-subject sleep-wake design. Twenty-eight healthy young adults (14 female; mean age 23.4 ± 3.1 years) encoded scene-label associations and were tested immediately and after either a 90-min daytime nap or a matched wake interval. During retrieval, learned and new scenes were presented at five levels of visual completeness. A separate pre-encoding baseline nap assessed individual sleep physiology without prior learning.

Sleep did not generally improve performance across all retrieval conditions. Instead, it selectively enhanced consolidation of learned scenes when visual cues were maximally degraded (p = .001) indicating increased pattern completion. No corresponding sleep effect was found for new scenes (all p > .31), suggesting that the benefit was specific to the recovery of previously encoded scene representations. Slow-wave amplitude during the post-encoding nap predicted consolidation of learned scenes under the most degraded condition (p = .011; FDR-corrected p = .042), whereas baseline slow-wave amplitude did not (p > .11).

These findings suggest that post-encoding sleep facilitates cue-based recovery of learned visual representations from strongly degraded input, and that this benefit is linked to slow-wave amplitude during post-encoding sleep. Together, these results link sleep-dependent consolidation to visual pattern-completion-like retrieval and extend previous work on sleep-related memory transformation from verbal and relational paradigms to the recovery of learned scene representations from degraded cues.

## Introduction

Episodic memory often requires the recovery of complete memory representations from partial or degraded cues. This process, commonly referred to as pattern completion, is considered a core hippocampal computation and has been linked to autoassociative dynamics within CA3 networks (Marr, 1971; McNaughton & Morris, 1987; Nakazawa et al., 2002; Neunuebel & Knierim, 2014; Treves & Rolls, 1994). When only a subset of the original input pattern is presented, CA3 recurrent networks can reinstate the complete stored representation, enabling recognition from minimal cues (Guzman et al., 2016). In humans, fMRI studies have confirmed that hippocampal activity during retrieval reflects the reinstatement of complete event representations from partial cues, with holistic pattern completion subsequently localized specifically to CA3 by ultra-high-field 7T fMRI (Grande et al., 2019; Horner et al., 2015).

Pattern completion is functionally complemented by *pattern separation* in the dentate gyrus (DG), which generates sparse, orthogonalized representations that reduce interference between similar memories (Hunsaker & Kesner, 2013; Rolls, 2016). Recent work has revised the simple DG-separates/CA3-completes framework: both computations are graded, depend on input similarity and task demands, and are embedded in broader cortico-hippocampal networks including prefrontal and parietal corteices (Amer & Davachi, 2023; Deuker et al., 2014; Liu et al., 2016). This graded architecture allows episodic memory to remain both specific and flexibly accessible depending on the degree of cue overlap at retrieval (McClelland et al., 1995; Yassa & Stark, 2011).

Sleep plays a central role in memory consolidation, with non-rapid eye movement (NREM) sleep serving as a critical window for hippocampus-dependent processing (Klinzing et al., 2019; Rasch & Born, 2013). According to the active systems consolidation hypothesis, slow oscillations (SOs) during NREM sleep orchestrate thalamocortical spindles and hippocampal sharp-wave ripples (SWRs) in a temporally nested hierarchy: SO upstates drive spindle bursts, which in turn cluster hippocampal ripples in their troughs, creating precisely timed frames for hippocampal reactivation and neocortical transfer (Diekelmann & Born, 2010; Rasch & Born, 2013; Staresina et al., 2015). Memory consolidation during sleep critically depends on this temporal coordination, and disrupting SO-spindle coupling impairs subsequent memory for associative and relational information (Latchoumane et al., 2017). Complementary learning systems (CLS) models predict that NREM-driven reactivations allow the hippocampus to train neocortical networks on shared representational structure, thereby strengthening neocortical representations that supports flexible memory retrieval, including completion from partial cues (Kumaran et al., 2016; McClelland et al., 1995).

A first line of evidence that sleep shapes hippocampal pattern-processing computations came from our earlier work using the Mnemonic Similarity Task (MST). Using the MST, we previously demonstrated that a night of sleep stabilizes mnemonic discrimination, with sleep-related improvements changes associated with NREM oscillatory markers (i.e., spindle and SO density; Hanert et al., 2017). Subsequent translational work in LGI1-LE patients using both the MST and the Virtual Morris Water Maze confirmed that DG/CA3 subfield integrity is necessary for sleep-dependent pattern separation, while residual CA3 function supports pattern completion even in a structurally compromised hippocampus (Rave et al., 2026).

Direct evidence that sleep facilitates pattern *completion* has begun to emerge. Recent work by Lutz et al. (2024) demonstrated that sleep supports pattern completion in multielement event memories by strengthening weak and indirect associations and preserving the co-retrieval of multiple event elements from a single cue. Another study (Joensen et al., 2024) used fMRI to show that hippocampal pattern completion signals remain robust following a 24-hour interval containing sleep, with holistic neocortical reinstatement emerging alongside persistent hippocampal involvement, indicating that consolidation redistributes rather than replaces the hippocampal basis of completion.

Despite this progress, critical gaps remain. Prior human studies employed verbal associative paradigms (Lutz et al., 2024) or inferred pattern completion indirectly from MST performance (Hanert et al., 2017), without directly measuring the retrieval of stored visual memory representations from parametrically degraded visual inputs. Most protocols used overnight sleep, which conflates sleep effects with circadian influences on encoding and retrieval. Finally, no prior study has included a pre-encoding baseline sleep recording to distinguish encoding-specific (state-dependent) effects from trait-like individual differences in sleep physiology, leaving open whether associations between sleep electroencephalogram (EEG) measures and memory performance reflect active, encoding-specific memory reprocessing or stable individual differences in sleep physiology.

To address these questions, we combined a within-subject sleep–wake design with the Memory Image Completion task, which presents learned and new scenes at different levels of visual completeness (Vieweg et al., 2019; Vieweg et al., 2015). This task allows pattern-completion-like retrieval to be assessed directly by testing whether learned scenes can be recognized when only limited visual information is available. In addition, participants completed a baseline nap before the sleep condition, allowing us to distinguish post-encoding sleep physiology from stable individual differences in sleep architecture. We hypothesized that a post-encoding nap would selectively enhance consolidation of learned scenes under conditions of maximal cue degradation. We further predicted that this benefit would be associated with NREM physiological markers of sleep-dependent memory consolidation during the post-encoding nap, consistent with sleep-dependent reprocessing of encoded scene representations.

## Methods

### Participants

Twenty-eight healthy young adults participated in the study (14 female, mean age = 23.36 years, SD = 3.12, range = 18–30 years). The average interval between experimental conditions (sleep vs. wake) was 39.04 days (SD = 23.68, range = 19–138 days). One participant (male) was excluded from analyses involving sleep parameters due to poor-quality sleep recordings but was retained for behavioral analyses. Participants were recruited via student mailing lists at the University of Kiel and received monetary compensation. None reported neurological or psychiatric disorders or regular use of medication (except oral contraceptives). Participants were instructed to refrain from caffeine intake after noon, alcohol consumption, and daytime napping on experimental days. All provided written informed consent. The study was approved by the local ethics committee of the University of Kiel and conducted in accordance with the Declaration of Helsinki.

An a priori power analysis was conducted for the primary simple linear regression model examining the association between post-encoding nap sleep parameters and memory consolidation. Assuming a large effect size (Cohen’s f²=0.35), an alpha level of .05, and a desired power of .80, the required sample size was estimated at N = 25. The final sample of N = 28 was therefore powered primarily to detect large effects. Multiple regression models including baseline nap values were treated as secondary analyses.

### Experimental design and procedure

The study used a within-subject design comprising a sleep condition and a wake condition. The order of the two conditions was counterbalanced across participants. All sessions were conducted in the sleep laboratory of the Department of Neurology at the University of Kiel. Participants completed a separate baseline nap session on the day preceding the sleep condition. The baseline nap was recorded using polysomnography and served to assess individual sleep physiology in the absence of preceding learning. In both experimental conditions, participants completed the Memory Image Completion task, consisting of an encoding phase, an immediate retrieval test, and a delayed retrieval test, seperated by a retention interval. In the sleep condition, the retention interval consisted of a 90-min post-encoding nap with polysomnographic (PSG) recording. Delayed retrieval testing started approximately 30 min after awakening. In the wake condition, participants completed the identical encoding and retrieval protocol but remained awake during the retention interval, which was matched in duration to the corresponding sleep interval. During the wake interval, participants watched neutral nature documentaries. Parallel versions of the MIC task were used across conditions, and the two experimental sessions were separated by at least two weeks to minimize proactive interference and repeated-exposure effects.

### Memory Image Completion Task

The Memory Image Completion (MIC) task was used to assess pattern completion and pattern separation and was adapted from the original paradigm developed by Vieweg et al. (2019; 2015). The task consisted of black-and-white line drawings of indoor scenes, each associated with a semantic room label (e.g, kitchen). For the present sleep study, the paradigm was modified to include four structurally equivalent parallel stimulus sets, allowing independent task versions across the sleep and wake conditions and across immediate and delayed retrieval. Each set comprised four target scenes to be learned, four additional scenes used during the learning-criterion phase, and four new foil scenes used during retrieval. Within each experimental condition, participants learned two sets of four target scenes, resulting in eight learned scenes per condition.

During encoding, participants learned the association between each target scene and its corresponding semantic room label. On each learning trial, the room label was presented first, followed by the corresponding scene, and all scene-label pairs were presented repeatedly in randomized order. Encoding was followed by a criterion phase to ensure successful acquisition of the scene-label associations. In this phase, the four learned target scenes were presented intermixed with four unlearned scenes. For each scene, participants first indicated whether it had been presented during encoding and, if so, selected the corresponding room label from a list. Feedback was provided after each response. Participants proceeded to the retrieval phase only after each learned target scene had been correctly identified and labelled on three consecutive presentations.

During retrieval, four learned target scenes and four new foil scenes from the respective stimulus set were presented at five levels of visual completeness (100%, 35%, 21%, 12%, and 5%). Lower completeness levels were created by progressively masking the scenes with white circles, resulting in increasingly degraded retrieval cues. Each retrieval phase therefore comprised 40 stimuli, consisting of four learned and four new foil scenes across five completeness levels, which were presented in randomized order. On each trial, participants selected the label corresponding to one of the learned scenes or indicated that none of the learned scenes was present, followed by a confidence rating (1 not at all confident, 5 very confident). Correct identification of learned target scenes was used as an index of pattern completion, reflecting retrieval of a learned representation from partial visual input. Correct rejection of new foil scenes was used as an index of pattern separation, reflecting the ability to distinguish novel scenes from previously learned representations. In addition, a pattern completion bias score was calculated as learned accuracy minus new accuracy, with positive values indicating a stronger bias toward completion of learned representations and negative values indicating a stronger bias toward separation of new stimuli. Consolidation scores were calculated as delayed minus immediate retrieval performance.

### Polysomnography and sleep EEG analysis

Sleep parameters were selected a priori based on their theoretical relevance for memory consolidation. We focused on four complementary measures capturing distinct aspects of sleep physiology: slow-wave amplitude as an index of SO strength, spindle density during NREM sleep as a commonly used marker of spindle occurrence, spindle amplitude as an index of spindle strength, and slow oscillation-spindle coupling, quantified using normalized direct phase-amplitude coupling, reflecting the extent to which spindle amplitude is systematically modulated by the phase of the SO.

Polysomnography was recorded using the SOMNOscreen EEG 10–20 system (Somnomedics, Germany). EEG signals were acquired from F3, F4, C3, C4, P3, P4, O1, and O2 according to the international 10-20 system, referenced to linked mastoids, with AFz serving as ground. Electrooculography (EOG) was recorded from electrodes placed around the eyes, and electrocardiography (ECG) was additionally acquired. EEG signals were digitized at 256 Hz and band-pass filtered using between 0.2 and 30 Hz; EOG signals between 0.2 and 10 Hz; and ECG signals with a 50 Hz low-pass filter.

Sleep stages were scored offline by a trained rater according to the criteria of the American Academy of Sleep Medicine. For each participant, total sleep time and time spent in N1, N2, N3, and rapid eye movement (REM) sleep were determined. Sleep onset was defined as the first occurrence of N1 followed by N2 sleep after lights off. Movement artifacts and visually identified EEG artifacts were excluded from subsequent analyses. Heavily artefact-contaminated channels were excluded from further analyses.

Quantitative sleep EEG analyses were performed in Python using YASA version 0.6.5. Analyses were restricted to artifact-free NREM sleep, defined as N2 and N3 sleep stages. All sleep EEG parameters were first extracted separately for each participant and channel and subsequently averaged across all EEG channels to obtain one value per participant if not otherwise stated.

### Slow-wave detection and slow-wave-spindle coupling

Slow waves were detected using the sw_detect function implemented in YASA. Detection was performed during NREM sleep in the frequency range of 0.3-1.5 Hz. Briefly, EEG signals were band-pass filtered in the slow-wave range using a finite impulse response filter. Candidate slow waves were identified based on negative trough amplitudes between -40 and -200 µV and positive peaks between 10 and 150 µV. Events were retained if their peak-to-peak amplitude was between 60 and 400 µV, the duration of the negative deflection was between 0.3 and 1.5 s, and the duration of the positive deflection was between 0.1 and 1.0 s. For each detected slow wave, peak-to-peak amplitude was extracted as the amplitude difference between the positive peak and the negative trough. The primary slow-wave outcome was mean slow-wave peak-to-peak amplitude. Values were averaged for all detected slow waves across all channels for each participant.

Slow-wave-spindle coupling was quantified using normalized direct phase-amplitude coupling implemented within YASA’s sw_detect function. Coupling was calculated between the phase of the slow-wave-filtered signal and the amplitude of sigma-band activity in the frequency range of 12-15 Hz. The coupling window was centered on the negative slow-wave peak and extended ±1 s around the trough. For each slow wave, the timing of the maximum sigma amplitude, the slow-wave phase at this sigma maximum, and the normalized direct phase-amplitude coupling value were extracted. The primary coupling outcome was mean normalized direct phase-amplitude coupling, averaged across slow waves within each channel and subsequently across channels for each participant.

### Sleep spindle detection

Previous to the spindle detection, the power spectrum was fitted between 1-30 Hz using the FOOOF toolbox in Python. The two individual spindle frequency peaks were extracted within the 8-16 Hz range. The center frequency of each peak was then used to detect the sleep spindles using the spindles_detect function implemented in YASA. Detection was performed during NREM sleep in the spindle frequency range of 9-11 Hz for slow spindles and 12–15 Hz for fast spindles. The broadband reference range was set to 1-30 Hz. Candidate spindle events were identified based on a combination of sigma-band relative power, moving correlation between the broadband and sigma-filtered signal, and root mean square amplitude of the sigma-filtered signal using YASA’s spindles_detect function. Events were retained if their duration was between 0.5 and 2.0 s. Spindles occurring within 1000 ms of each other were merged.

For each participant and channel, spindle density and spindle amplitude were extracted. Spindle density was defined as the number of detected spindles normalized to the number of 30-s NREM (N2 and N3) epochs. Spindle amplitude was defined as peak-to-peak amplitude of the detrended spindle in the broadband-filtered signal. The primary spindle outcomes were mean spindle density and mean spindle amplitude. Both variables were first calculated separately for each channel and then averaged across all channels for each participant.

### Statistical analyses

All analyses were conducted in Python (v3.13) using *pandas*, *statsmodels*, *scipy*, and *seaborn*. The statistical analyses of the sleep architecture were conducted in R (v4.5.1) using *dplyr* and *tidyr* packages Macro sleep architecture measures and a priori selected sleep physiology measures were compared between the baseline nap and the post-encoding nap using paired-samples t-tests or Wilcoxon signed-rank tests, depending on whether the data were normally distributed, as assessed with Shapiro–Wilk tests. Because these comparisons involved a small, predefined set of related sleep variables and served to characterize differences between the two naps, p-values were evaluated using the Holm-Bonferroni procedure to control the family-wise error rate. For the sleep-memory regression analyses, which involved multiple exploratory associations between sleep parameters and behavioral outcomes, we controlled the false discovery rate using the Benjamini-Hochberg procedure.

Consolidation scores were calculated as delayed minus immediate recall performance separately for condition, stimulus type, and completeness level. The primary dependent variables were learned-stimulus accuracy, new-stimulus accuracy, and a bias score defined as learned minus new accuracy, with higher values indicating stronger pattern completion. For each dependent variable, linear mixed-effects models were fitted with condition (sleep, wake), stimulus completeness (100%, 35%, 21%, 12%, 5%), and their interaction as fixed effects. Completeness was modeled categorically, and subject was included as a random intercept. Models were estimated using maximum likelihood. Fixed effects are reported as β estimates with standard errors, z-values, p-values, and 95% confidence intervals.

Model assumptions were assessed using residual histograms, Q-Q plots, residuals-versus-fitted plots, and tests of normality and homoscedasticity. Because random-intercept variance was close to zero in some diagnostic fits, assumption checks were additionally inspected using fixed-effects models with the same predictor structure. Robustness was assessed by repeating analyses after excluding observations with standardized residuals exceeding ±3.

Associations between sleep physiology and memory performance were examined using linear regression models at the 5% completeness level, where the behavioral analyses revealed sleep-related effects for learned accuracy and pattern completion bias. For each outcome, separate models were fitted for each sleep parameter, with the test-nap parameter entered as predictor and the corresponding baseline parameter as covariate. Sleep predictors were z-standardized. p-values were FDR-corrected separately within each behavioral outcome across the primary sleep predictors. Three sensitivity models complemented the primary analysis: (1) test-nap only, (2) baseline only, and (3) test-minus-baseline difference scores. The prediction that test-nap but not baseline slow-wave amplitude would predict memory performance was specified a priori as the key evidence for encoding-specific reprocessing. Assumptions for the sleep-memory regression models were evaluated using residual plots, studentized residuals, leverage, Cook’s distance, variance inflation factors, and correlations between baseline and test-nap predictors. Models with highly correlated baseline and test-nap spindle measures were interpreted cautiously. All tests were two-tailed with α = .05.

## Results

To investigate the effects of sleep on memory consolidation and pattern completion, linear mixed-effects models were fitted on the consolidation scores (delayed minus immediate recall) for learned stimuli, new stimuli, and the bias score (learned minus new stimuli), with condition (sleep vs. wake) and stimulus completeness (100%, 35%, 21%, 12%, 5%) as fixed effects and subject as a random intercept (Figure 2). Performance approached ceiling at high completeness levels and declined as stimulus completeness decreased, with the largest variability observed at the 5% level. Table S1 provides descriptive statistics for learned stimuli, new stimuli, and the bias measure across completeness levels before and after the sleep and wake retention intervals.

**Figure 1.**
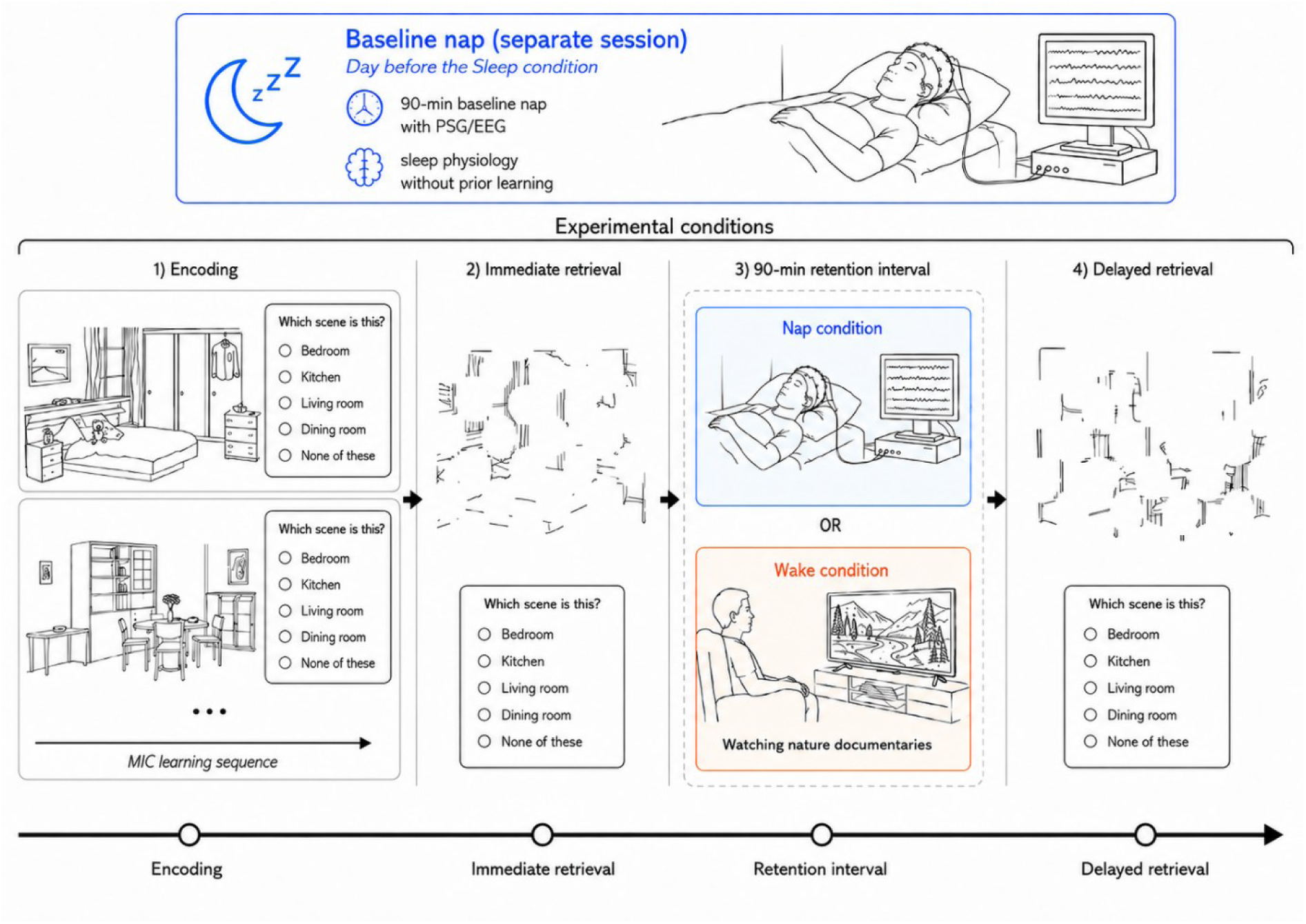
Experimental design. The study followed a within-subject design with counterbalanced sleep and wake conditions. Participants completed a separate baseline nap session on the day preceding the sleep condition to assess sleep physiology during a 90-min daytime nap without prior learning. In both experimental conditions, the task protocol comprised encoding, immediate retrieval, a 90-min retention interval, and delayed retrieval. In the sleep condition, participants took a 90-min post-encoding nap with simultaneous PSG/EEG recording, and delayed retrieval started approximately 30 min after awakening. In the wake condition, participants remained awake for an equivalent interval while watching neutral nature documentaries. Parallel versions of the MIC were used across conditions. The two experimental sessions were separated by at least two weeks.

**Figure 2.**
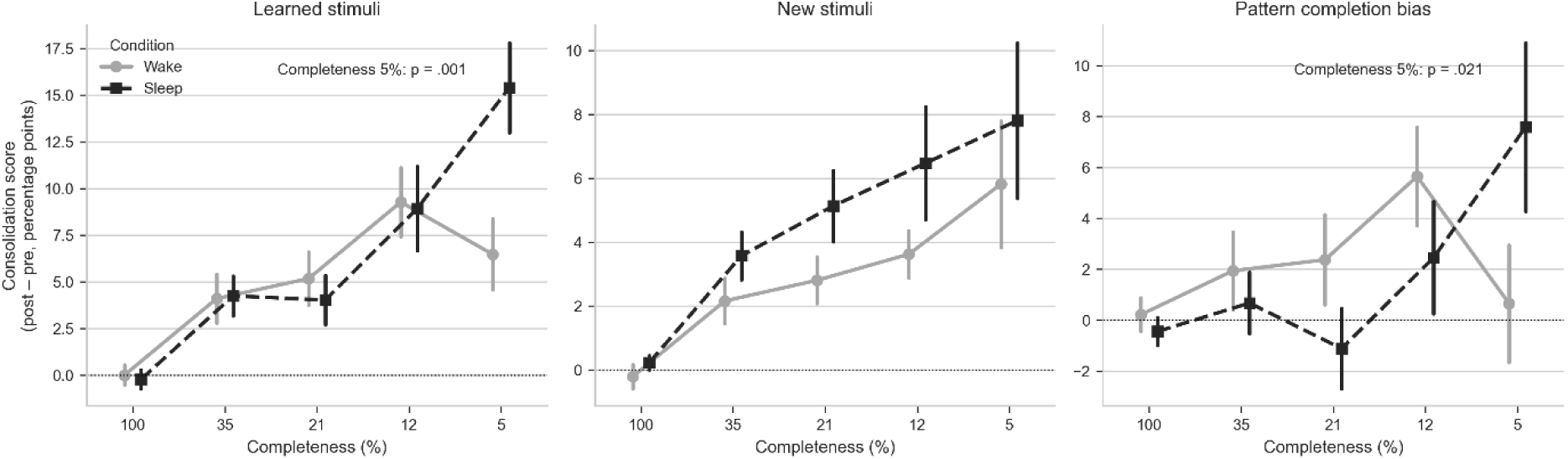
Consolidation scores across stimulus completeness levels for learned stimuli, new stimuli, and pattern completion bias. Consolidation scores were calculated as changes from immediate to delayed recall in accuracy or bias and are shown separately for the sleep and wake conditions. Points indicate group means, and error bars represent the standard error of the mean. Significant sleep-related effects were observed only for learned stimuli and the pattern completion bias at the 5% completeness level.

### Pattern Completion Performance (learned stimuli)

For learned stimuli, there was no main effect of condition, indicating that overall consolidation did not differ between the sleep and the wake condition (β = -0.002, SE = 0.019, z = -0.10, p = .919, 95% CI [- 0.040, 0.036]). However, a significant main effect of stimulus completeness was observed, with greater consolidation scores at lower levels of stimulus completeness (35%: β = 0.041, p = .031, 95% CI [0.004, 0.078]; 21%: β = 0.052, p = .007, 95% CI [0.014, 0.089]; 12%: β = 0.093, p < .001, 95% CI [0.055, 0.130]; 5%: β = 0.065, p = .001, 95% CI [0.027, 0.102]). A significant interaction between condition and stimulus completeness was found at the lowest completeness level (5%), indicating greater consolidation following sleep compared to wake (β = 0.092, SE = 0.027, z = 3.38, p = .001, 95% CI [0.038, 0.145]). No interaction effects were observed at higher completeness levels (all p > .70).

### Pattern Separation Performance (new stimuli)

For new stimuli, there was no main effect of condition (β = 0.004, SE = 0.017, z = 0.26, p = .798, 95% CI [-0.029, 0.038]). A main effect of stimulus completeness was observed, with greater immediate to delayed recall changes at lower levels of stimulus completeness, reaching significance at 12% (β = 0.038, p = .023, 95% CI [0.005, 0.071]) and 5% completeness (β = 0.060, p < .001, 95% CI [0.027, 0.093]). No interaction between condition and stimulus completeness was observed (all p > .31), indicating that sleep did not significantly affect performance changes for new stimuli, arguing against a general criterion shift as the explanation for the sleep benefit observed for learned stimuli.

### Pattern Completion Bias (learned – new stimuli)

For the bias measure, there was no main effect of condition (β = -0.006, SE = 0.023, z = -0.27, p = .786, 95% CI [-0.052, 0.039]). A significant main effect of stimulus completeness was observed at 12% completeness (β = 0.054, p = .018, 95% CI [0.009, 0.099]). A significant interaction between condition and stimulus completeness emerged at the lowest completeness level (5%), indicating a greater increase in pattern completion bias following sleep compared to wake (β = 0.076, SE = 0.033, z = 2.31, p = .021, 95% CI [0.012, 0.140]). No interaction effects were found at higher completeness levels (all p > .39).

### Sensitivity analysis

To assess robustness, all models were re-estimated after excluding observations with standardized residuals exceeding ±3. The critical interaction between condition and stimulus completeness at the 5% level remained significant for learned stimuli (β = 0.083, p = .007) and for the bias measure (β = 0.084, p = .016), whereas no such interaction emerged for new stimuli (β = -0.010, p = .639). Thus, the main pattern of results was not driven by a small number of outlying observations. Model diagnostics indicated small to moderate deviations from normality and homoscedasticity, particularly at the lowest completeness level, where variability was highest. Because random intercept variance in the consolidation-score models was close to zero, diagnostic plots were additionally inspected using corresponding fixed-effects models. Overall, no violations were considered severe enough to invalidate interpretation of the main effects.

### Sleep architecture and electrophysiological characteristics

We compared sleep architecture between the baseline nap and the post-encoding test nap. After Holm-Bonferroni correction for multiple comparisons, only total sleep time remained significant, with participants sleeping longer during the post-learning test nap than during the baseline nap. No other sleep architecture measures differed significantly (Table 1).

**Table 1.** Sleep stages in minutes and percent (M ± SEM) for the baseline and post-encoding test naps.

|  | baseline (n = 27) | test (n = 27) | t(26)/ W | p |
| --- | --- | --- | --- | --- |
| TiB (min) | 95.48 ± 0.73 | 99.48 ± 2.69 | 154 | 0.59<sup>#</sup> |
| TST (min) | 63.67 ± 3.84 | 76.62 ± 4.66 | -4.01 | **0.0005<sup>a</sup>** |
| N1 (min) | 4.79 ± 1.12 | 3.27 ± 0.55 | 138 | 0.72<sup>#</sup> |
| N2 (min) | 30.77 ± 2.85 | 38.33 ± 3.32 | -2.61 | **0.015** |
| N3 (min) | 15.73 ± 3.52 | 19.46 ± 3.09 | -1.11 | 0.28 |
| REM (min) | 12.38 ± 2.2 | 15.56 ± 2.47 | -1.20 | 0.24 |
| WASO (min) | 9.65 ± 1.96 | 6.08 ± 2.17 | 251.5 | **0.017<sup>#</sup>** |
| N1 (%) | 7.97 ± 2.02 | 4.78 ± 0.87 | 190 | 0.47 |
| N2 (%) | 53.21 ± 5.01 | 51.13 ± 3.19 | 0.51 | 0.62 |
| N3 (%) | 21.8 ± 4.88 | 24.79 ± 4.25 | -0.73 | 0.47 |
| REM (%) | 17.02 ± 2.88 | 19.3 ± 2.9 | -0.63 | 0.53 |
| WASO (%) | 0.21 ± 0.06 | 0.23 ± 0.18 | 254 | **0.014<sup>#</sup>** |

TiB, time in bed; TST, total sleep time; N1, sleep stage 1; N2, sleep stage 2; N3, sleep stage 3; REM, rapid eye movement sleep, WASO, wake after sleep onset. Percentages are expressed relative to TST. Differences between baseline and post-encoding test nap were tested using paired-samples t-tests or Wilcoxon signed-rank tests (#), depending on data distribution. <sup>a</sup> Significant after Holm-Bonferroni correction for multiple comparisons.

We next compared a priori selected electrophysiological sleep measures between the baseline nap and the post-encoding nap, including SO density, SO peak-to-peak amplitude, spindle density, spindle amplitude, and SO-spindle coupling indexed by ndPAC. None of these measures remained significant after Holm-Bonferroni correction for multiple comparisons. Thus, the baseline and post-learning naps did not differ significantly in SO, spindle, and coupling characteristics (Table 2).

**Table 2.**
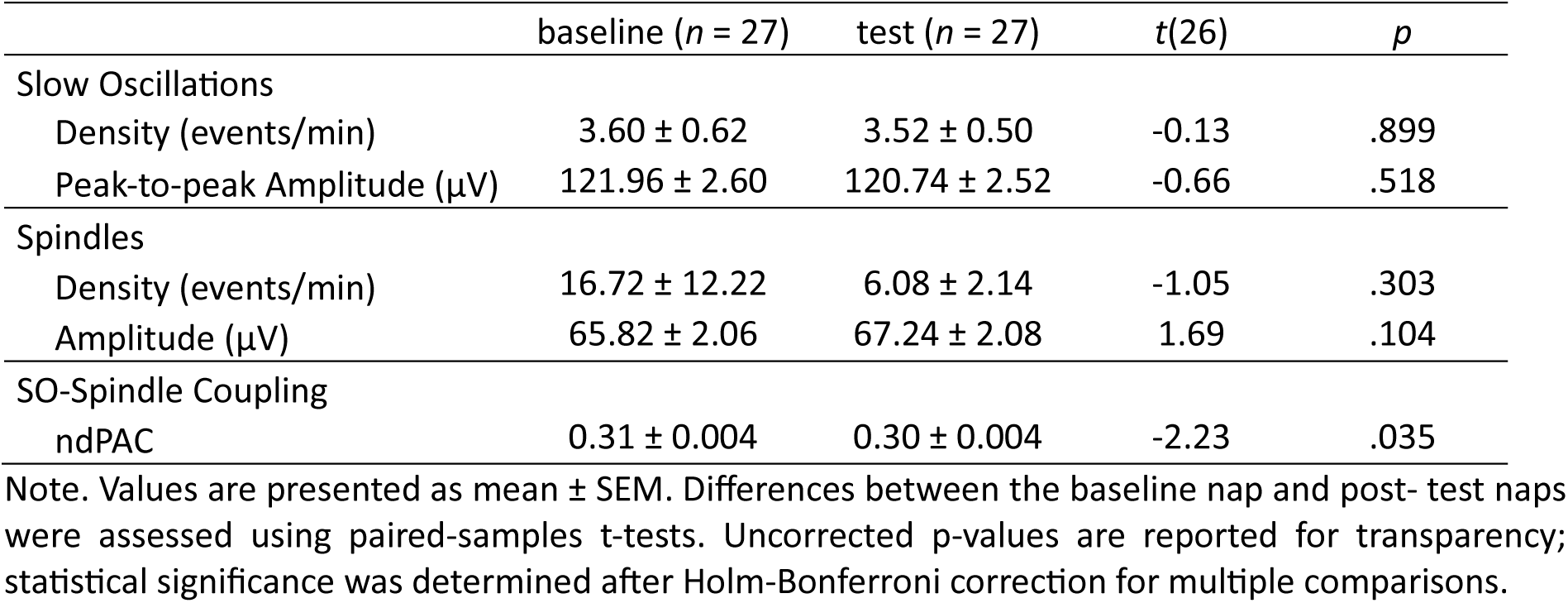
Slow oscillation and spindle characteristics during the baseline and post-encoding test naps.

### Slow-Wave amplitude during the post-encoding nap predicts consolidation

Given that the behavioral analyses revealed sleep-related improvements only at the lowest level of stimulus completeness (5%) for learned stimuli and pattern completion bias, subsequent analyses focused on this condition to examine whether inter-individual differences in sleep physiology were associated with these effects. Accordingly, separate linear regression models were computed for learned accuracy and bias at the 5% completeness level, including post-encoding test sleep parameters while controlling for the corresponding baseline measures. For learned accuracy, greater slow-wave amplitude during the post-encoding test nap significantly predicted consolidation performance when controlling for baseline slow-wave amplitude (β = 0.090, SE = 0.035, z = 2.56, p = .011, 95% CI [0.021, 0.159]). This effect remained significant after correction for multiple comparisons using the false discovery rate (p = .042). No other sleep parameter significantly predicted learned accuracy after correction (all p > .50). Baseline sleep parameters were not significant predictors (all p > .11). For the pattern completion bias, no significant associations with any sleep parameter were observed (all p’s > .80). Together, these findings suggest that slow-wave amplitude during the experimental nap was selectively associated with improved consolidation of learned stimuli under maximal pattern-completion demands.

To assess whether the observed association was specifically related to sleep during the experimental nap, additional models were computed using test-nap parameters alone, baseline parameters alone, and test-minus-baseline difference scores. For learned accuracy, the association with slow-wave amplitude was also significant in the test-only model (β = 0.049, SE = 0.025, z = 2.01, p = .045, 95% CI [0.001, 0.097]) and in the test-minus-baseline model (β = 0.051, SE = 0.024, z = 2.11, p = .035, 95% CI [0.004, 0.099]). In contrast, baseline-only models did not yield significant effects (see Figure 3 A and B). Across the primary model and the three sensitivity analyses, the observed pattern provides converging evidence from complementary analytical approaches that the association between slow-wave amplitude and memory is linked to post-encoding sleep rather than reflecting a stable trait. Importantly, the baseline nap was administered in the same setting, at the same time of day, with the same EEG montage, differing only in the absence of to-be-consolidated material. This design allowed us to test whether individual slow-wave amplitude predicts pattern completion independently of memory encoding. The absence of a significant association in the baseline-only model argues against this interpretation.

**Figure 3.**
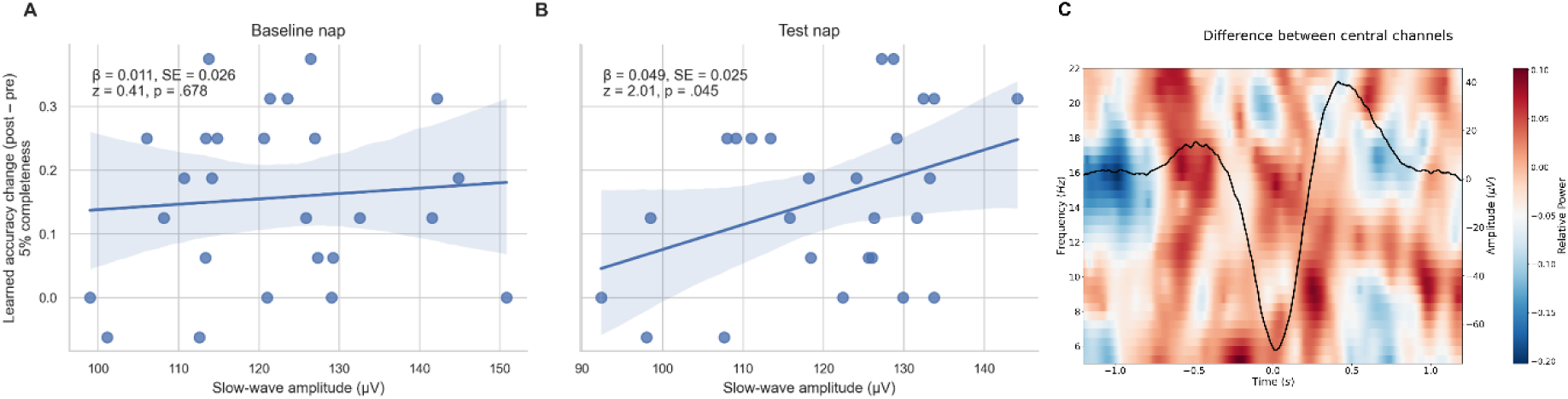
Sleep physiology during the baseline and post-encoding test naps and its association with learned-stimulus consolidation. (A, B) Association between slow-wave amplitude and learned-stimulus consolidation at 5% completeness. Baseline slow-wave peak-to-peak amplitude (A) and post-encoding test-nap slow-wave peak-to-peak amplitude (B) are plotted against immediate to delayed recall changes in learned accuracy. Points represent individual participants, and lines indicate fitted regressions with 95% confidence intervals. Regression lines correspond to the baseline-only and post-encoding-test-nap-only models, respectively. Only test nap slow-wave amplitude was significantly associated with learned-stimulus consolidation. (C) Time-frequency representation of differences between baseline and test nap over central channels. Warm colors indicate higher relative power in the test nap compared with baseline, whereas cool colors indicate lower relative power in the test nap compared with baseline. The black line depicts the average EEG waveform across the same central channels and is shown on the right y-axis. No significant clusters were detected using cluster-based permutation testing.

## Discussion

The present study provides three main findings. First, we show that a daytime nap selectively enhanced the recognition of learned visual images at the lowest level of stimulus completeness (5%), where successful retrieval requires reconstructing a stored representation from maximally sparse input. This effect was specific to previously learned stimuli and constituted an increase in pattern completion bias. Second, individual differences in slow-wave amplitude during the post-encoding test nap predicted this selective benefit, even after controlling for baseline slow-wave amplitude indexed by a baseline nap. Third, baseline slow-wave amplitude alone did not predict performance, suggesting that the association is related to consolidation-specific rather than trait-level slow-wave activity. Together, these findings indicate that NREM slow-wave activity during post-encoding sleep facilitates the stabilization and flexible reinstatement of visual memory representations, thereby enhancing memory-based pattern completion from sparse cues.

### Selectivity and theoretical significance of the 5% completeness effect

Our findings are consistent with the growing view that sleep-dependent memory consolidation does not merely preserve individual memories but can enhance the accessibility, robustness, and relational organization of memory representations. Within the framework of active systems consolidation, newly encoded hippocampal memories are reactivated during post-learning sleep and gradually redistributed within hippocampal-neocortical networks. Such reprocessing is thought to render memory traces more resistant to interference and more reliably accessible under high retrieval demands. Previous work on sleep and hippocampal computations has primarily focused on mnemonic discrimination and pattern separation (Hanert et al., 2017). Similarly, Doxey et al. (2018) reported sleep-related effects on memory specificity and neural response patterns associated with pattern separation. More recently, Lutz et al. (2024) extended this literature to pattern completion by showing that sleep strengthens the associative structure of multielement event memories and improves the ability to retrieve multiple event elements from a single cue. Together, these findings suggest that sleep can support both the distinctiveness of similar memories and the flexible recovery of complete representations from partial input.

An important methodological strength of the present study is that pattern completion was assessed more directly than in previous sleep studies using mnemonic discrimination paradigms. The MST used in earlier work is well established as a measure of mnemonic discrimination and pattern separation, because participants are required to distinguish studied items from highly similar lures and novel foils (Stark et al., 2013). However, pattern completion is typically inferred indirectly in this task, for example from false recognition of similar lures as old items or from neural response profiles interpreted as completion-like (Liu et al., 2016). In contrast, the Memory Image Completion task was specifically designed to assess pattern-completion-based retrieval by presenting learned scenes at parametrically reduced levels of visual completeness (Vieweg et al., 2019; Vieweg et al., 2015). This task therefore allows direct assessment of whether a stored representation can be recovered from partial or degraded cues, closely matching the classical definition of pattern completion. Accordingly, the present findings complement previous MST-based sleep studies by showing that sleep can facilitate not only the preservation of memory specificity (Doxey et al., 2018; Hanert et al., 2017), but also the retrieval of learned scene representations from highly degraded visual input.

The restriction of the sleep benefit to the 5% completeness level can be interpreted in at least two complementary ways. First, performance at higher completeness levels was close to ceiling, particularly at 100%, 35%, and 21% completeness (see Table S1). Thus, these conditions may have provided limited sensitivity to detect additional sleep-related improvements. The absence of sleep effects at higher completeness levels should therefore not be interpreted as evidence against sleep-dependent consolidation, because sufficient visual information may have allowed accurate recognition already before the retention interval. Second, the selectivity of the effect is theoretically coherent with the computational demands of pattern completion. At higher completeness levels, recognition may be supported by perceptual matching or by the use of diagnostic visual features, without requiring full reinstatement of the stored representation. At 5% completeness, by contrast, the available sensory information is so sparse that successful recognition is likely to rely more strongly on the ability to reinstate a complete stored representation from minimal input. In computational terms, this corresponds to the operation attributed to hippocampal autoassociative networks, particularly CA3, where partial input can trigger convergence toward a stored attractor state (Marr, 1971; Rolls, 2016; Treves & Rolls, 1994). Although the present behavioral data do not allow direct conclusions about CA3 mechanisms, they are consistent with the idea that sleep enhanced the internal accessibility or stability of learned representations, making their recovery from highly degraded cues more reliable.

Importantly, the absence of a corresponding sleep effect for new stimuli argues against a non-specific response bias or generalized response tendency account. If sleep had merely increased a tendency to endorse fragmented stimuli as learned, then new stimuli should also have been affected, particularly at low completeness levels. Instead, sleep selectively benefited learned stimuli and increased the pattern completion bias without impairing correct rejection of new stimuli. This dissociation suggests that sleep enhanced access to previously encoded scene representations rather than inducing a general completion tendency. Taken together, the present findings are consistent with a cue-based form of memory generalization: sleep strengthened the internal representation of learned scenes such that they could be reinstated from increasingly degraded sensory inputs. Pattern completion is the computational mechanism underlying this generalization capacity, because it allows a partial, degraded, or noisy retrieval cue to be mapped onto a stored representationand thereby enables recovery of the same memory across perceptually impoverished versions of the original input (Kumaran & McClelland, 2012; Marr, 1971; Norman & O’Reilly, 2003; Rolls, 2013; Treves & Rolls, 1994; Yassa & Stark, 2011). Thus, after sleep, learned representations could be retrieved from increasingly impoverished perceptual inputs, but this increased accessibility did not generalize indiscriminately to novel stimuli. One methodological consideration is that the use of a 90-minute daytime nap, rather than a full overnight sleep episode, may have limited the observable consolidation effects to conditions of particularly high retrieval demand. A nap may provide sufficient sleep-dependent processing to improve retrieval when task demands are particularly high, but may be too limited to produce broad improvements across easier retrieval conditions (Mednick et al., 2003). Prior nap studies using mnemonic discrimination tasks have not consistently shown sleep-related benefits, suggesting that a short sleep opportunity may be sufficient to reveal consolidation effects only under conditions of particularly high retrieval demand (Cellini et al., 2020; Reichardt et al., 2024). Thus, the present findings indicate that even a 90-min nap can enhance pattern-completion-like retrieval, but that this effect becomes behaviorally visible primarily when the retrieval cue is maximally degraded and the task strongly depends on the integrity of the consolidated representation.

### Slow-Wave amplitude as a specific marker of encoding-dependent reprocessing

The selective improvement after the post-encoding nap relative to the wake interval suggests that the observed consolidation benefit was not merely a consequence of time-dependent stabilization or repeated testing but was specifically linked to sleep occurring after learning. This behavioral benefit of the post-encoding nap is consistent with active systems consolidation accounts, according to which newly encoded hippocampus-dependent memories are reactivated during non-REM sleep and gradually redistributed within hippocampal-neocortical networks (Diekelmann & Born, 2010; Klinzing et al., 2019; Rasch & Born, 2013). A key role in this process has been attributed to SOs, which provide a temporal framework for coordinating thalamocortical spindles and hippocampal sharp-wave ripples, thereby aligning hippocampal replay with periods of increased cortical plasticity (Latchoumane et al., 2017; Staresina et al., 2015). The present physiological findings strengthen this interpretation. Slow-wave amplitude during the post-encoding nap predicted consolidation of learned stimuli at the lowest completeness level, even when individual baseline slow-wave amplitude was taken into account. Within this framework, greater slow-wave amplitude during the post-encoding test nap may reflect stronger synchronization of the SO upstate, potentially extending the temporal window that supports hippocampal SWRs and increasing the probability of successful memory reactivation and stabilization of recently encoded scene representations, rendering them more accessible from highly degraded retrieval cues. With respect to pattern completion, slow-oscillation-driven reactivation may reinforce the CA3 recurrent connectivity that underlies attractor dynamics: repeated replay of encoding-specific activity patterns during NREM may potentiate CA3 recurrent synapses through spike-timing-dependent plasticity, potentially deepening the attractor basin for each stored representation and thereby improving convergence from sparse cues at retrieval (Kumaran et al., 2016; Norman & O’Reilly, 2003).

A key strength of the present design is the inclusion of a baseline nap, which allowed us to distinguish post-encoding sleep physiology from stable individual differences in habitual sleep physiology. Slow-wave amplitude predicted consolidation of learned stimuli at 5% completeness during the post-encoding nap, even when baseline slow-wave amplitude was controlled, whereas baseline slow-wave amplitude alone showed no comparable association. This dissociation argues against a purely trait-based account, according to which individuals with generally higher slow-wave amplitude would be expected to exhibit better memory performance irrespective of whether encoding had occurred. Instead, the converging results of the primary model, the post-encoding test-nap-only model, and the test-minus-baseline model suggests that the observed association with slow-wave amplitude was specifically linked to sleep following encoding. Thus, the baseline-nap design supports the interpretation that post-encoding slow-wave amplitude reflects a learning-dependent sleep process that increased the accessibility or stability of newly encoded scene representations, enabling their later recovery from highly impoverished retrieval cues. This provides converging within-person evidence supporting a role of encoding-specific slow-wave modulation in memory consolidation.

The finding that spindle density, spindle amplitude, and SO-spindle coupling did not emerge as significant predictors may seem surprising given the prominent role of spindles in prior sleep-memory studies (e.g., Fernandez & Lüthi, 2020; Hanert et al., 2017; Klinzing et al., 2019; Rasch et al., 2007). Spindle mediated associative binding as proposed by Lutz et al. (2024) for multielement verbal associations may be most relevant when the consolidation task requires strengthening the inter-element links of a relational structure, a process particularly dependent on synaptic tagging and binding across distributed neocortical representations. By contrast, the consolidation of discrete visual scene representations that support holistic reinstatement from sparse cues may depend more strongly on the integrity of the slow-oscillatory cycle that supports memory reactivation. In this context, slow-wave amplitude may index the strength or synchrony of the slow-oscillation up- and down-state dynamics that gate periods of cortical excitability and coordinate the temporal nesting of spindles and hippocampal sharp-wave ripples (Clemens et al., 2007; Mölle et al., 2006). Consistent with this view, larger SO amplitudes and longer up-states have been linked to stronger overnight memory improvement, possibly because they provide a more reliable temporal window for hippocampal-neocortical communication and spindle-coupled plasticity (Heib et al., 2013). Moreover, recent work suggests that memory consolidation benefits not only from the presence of spindles, but from their precise coupling to the SO up-state, particularly within well-structured slow waves (Muehlroth et al., 2019; Ng et al., 2025; Niknazar et al., 2015). In this framework, the SO amplitude may have provided a more global marker of reactivation-supporting sleep physiology than aggregate measures of spindle density, spindle amplitude, or SO-spindle coupling. Thus, slow-wave amplitude may have provided a more upstream or global marker of reactivation-supporting sleep physiology than aggregate measures of spindle density, spindle amplitude, or SO-spindle coupling. Importantly, this interpretation does not imply that spindles are irrelevant for pattern-completion-related consolidation, but rather that the spindle measures used here may not have captured the specific physiological process most closely related to the recovery of discrete scene representations from highly impoverished cues.

### Pattern Completion, generalization, and the role of sleep

Sleep has long been proposed to do more than stabilize newly encoded information. Rather, repeated memory reactivation during sleep may reorganize memories, integrate them with existing knowledge, and gradually promote more abstract or gist-like representations (Inostroza & Born, 2013; Klinzing et al., 2019; Walker & Stickgold, 2010). Empirical support for this view comes from studies showing that sleep can facilitate relational inference and memory integration. For example, Ellenbogen et al. (2007) showed that sleep preferentially enhanced transitive inference for indirectly related item pairs, indicating that sleep can promote the flexible use of learned relations beyond directly encoded associations. Similarly, Werchan & Gómez (2013) reported that sleep enhanced transitive inference when learning was supported by reinforcement. However, evidence for sleep-dependent generalization is not uniform across paradigms. Studies using targeted memory reactivation have shown that sleep can preserve or weaken specific memory details without necessarily producing robust gains in generalization (Witkowski et al., 2021). Sleep-related memory transformation is therefore task-dependent, with the operative mechanism determined by the structure of the to-be-consolidated representation and the demands of subsequent retrieval. The present findings extent this literature by examining a more restricted form of generalization.

Generalization in the context of memory refers to the ability to apply stored knowledge to novel or degraded inputs, which is precisely what pattern completion enables when the available sensory input is sparse or distorted (Kumaran & McClelland, 2012; Zeithamova & Bowman, 2020). In contrast to studies of relational inference, category learning, or schema formation, the MIC task does not require participants to infer novel relationships across multiple memories or to extract shared regularities across events. Instead, successful performance requires retrieval of a specific learned scene from a degraded version of that same scene (Vieweg et al., 2019; Vieweg et al., 2015). The relevant form of flexibility is therefore best described as cue-based generalization: the ability to recover an encoded representation across perceptually different versions of the original input. This distinction is important for interpreting the relationship between the present findings and models of sleep-dependent abstraction. Lewis & Durrant (2011) proposed that overlapping replay during slow-wave sleep selectively strengthens shared elements across related memories, thereby promoting the emergence of gist and schema-like knowledge. In the present task, however, there was no need to extract common structure across multiple overlapping episodes. Instead, sleep appeared to improve the ability to recover an individual scene representation when the external cue provided minimal sensory information. Thus, the present effect may reflect a representation-specific, cue-based, representation-specific form of generalization rather than schema-level abstraction. This interpretation is also consistent with the selectivity of the behavioral effect. Sleep improved consolidation for learned stimuli at 5% completeness, but did not produce a corresponding effect for new stimuli. The sleep benefit therefore may have increased the range of cues from which learned representations could be successfully accessed, without inducing indiscriminate overgeneralization to novel scenes.

Finally, this interpretation connects the present findings to recent work on sleep and pattern completion. Lutz et al. (2024) showed that sleep strengthens weak and indirect associations within multielement event memories and improves the ability to retrieve multiple event elements from a single cue, suggesting that sleep can shape associative structures that support pattern completion. The present study extends this idea to visual scene completion. Together with the findings by Lutz et al. (2024), the present results suggest that sleep may support pattern completion through complementaty processes, both by strengthening associative links among multiple event elements and by increasing the accessibility of a specific scene representation from strongly degraded cues. Thus, across different paradigms, sleep may support flexible memory expression through different mechanisms. That is, associative integration when multiple event elements must be linked, schema-like abstraction when shared structure must be extracted across memories, and cue-based generalization when a specific representation must be recovered from partial input. The present study extends this idea to visual scene completion. Here, pattern completion was assessed as the recovery of a specific learned scene representation from parametrically degraded perceptual input. The observation that both forms of pattern completion appear to benefit from sleep, yet are associated with distinct oscillatory correlates (spindle activity vs. slow-wave amplitude), suggests that sleep may engage multiple complementary consolidation mechanisms depending on the nature and structure of the memory representation and retrieval demands.

### The DG-CA3 system: Pattern Separation, Pattern Completion, and sleep

The present findings may also be considered in the context of previous work suggesting that sleep supports complementary hippocampal computations involved in memory specificity and flexible retrieval. Prior MST-based work showed that sleep stabilizes mnemonic discrimination performance, consistent with a role of sleep in preserving the distinctiveness of overlapping memory representations (Doxey et al., 2018; Hanert et al., 2017). In contrast, the present MIC findings indicate that a post-encoding nap can facilitate recovery of learned scene representations from strongly degraded cues, consistent with enhanced pattern-completion-like retrieval. Together, these findings suggest that sleep supports both the separation of similar memories and the subsequent completion of learned representations, depending on task demands and the structure of the memory representation.

Within this framework, the present results may be viewed as complementary to both our previous MST-based work (Hanert et al., 2017) and the LGI1-LE findings reported by Rave et al. (2026). Pattern separation has been linked to the orthogonalization of overlapping inputs within DG/CA3 circuitry, whereas pattern completion has been associated with CA3 autoassociative dynamics and the reinstatement of stored representations from partial cues (J. K. Leutgeb et al., 2007; S. Leutgeb et al., 2004; Norman & O’Reilly, 2003; Yassa & Stark, 2011). Taken together, these findings are broadly consistent with the idea that sleep supports pattern separation and pattern completion through partly distinct NREM oscillatory mechanisms. (Hanert et al., 2017) found that overnight stabilization of lure discrimination was associated with spindle density, SO density, and SO-locked theta power, indicating that NREM oscillatory dynamics may contribute to the preservation of memory specificity. In the present study, by contrast, slow-wave amplitude during the post-encoding nap predicted consolidation of learned scene representations under maximal cue degradation. This pattern is consistent with the idea that stronger slow-oscillatory activity supports the reactivation or stabilization of representations needed for completion-based retrieval. However, because the present study did not include subfield-specific imaging or direct measures of hippocampal replay, the proposed DG-CA3 dissociation should be understood as a mechanistic hypothesis rather than a direct conclusion from the current data.

### Limitations and Future Directions

Several limitations warrant consideration. First, the sample size was modest and the study was primarily powered to detect large effects. Although the association between test-nap slow-wave amplitude and consolidation of learned stimuli was consistent across the primary model, the test-nap-only model, and the test-minus-baseline model, these findings should be interpreted cautiously and require replication in larger samples. Second, performance at higher completeness levels was close to ceiling, particularly for 100%, 35%, and 21% completeness. This likely reduced sensitivity to detect sleep-related benefits under easier retrieval conditions and may have contributed to the restriction of the behavioral effect to the 5% completeness level. However, ceiling effects cannot fully explain the observed pattern, as the sleep benefit was selective for learned stimuli and the bias measure, with no corresponding effect for new stimuli. Third, the study did not include neuroimaging or direct measures of hippocampal replay. Any interpretation in terms of hippocampal subfield mechanisms, including CA3-related pattern completion, therefore remains theoretical rather than directly demonstrated. Fourth, the use of a daytime nap limits generalization to overnight sleep, which includes multiple NREM-REM cycles and may support broader or more robust consolidation effects. Finally, slow-wave amplitude predicted consolidation of learned stimuli but not the pattern completion bias, despite the behavioral sleep effect in both outcomes. This suggests that the bias measure may capture additional cognitive components, such as response strategies or decision criteria, that are not directly indexed by slow-wave physiology. Future studies combining the MIC task with high-resolution hippocampal imaging, targeted memory reactivation, or closed-loop modulation of SOs will be important to test whether sleep-dependent improvements in completion-based retrieval are mediated by scene-specific reactivation and hippocampal subfield dynamics.

### Conclusions

A daytime nap selectively enhances retrieval of learned visual images from maximally sparse cues, indicating that post-encoding sleep supports cue-based recovery of stored representations under conditions of maximal perceptual degradation. The strength of NREM slow-wave activity during post-encoding sleep, but not during a pre-encoding baseline nap, predicts this benefit consistently across the primary model and three complementary sensitivity analyses, providing converging evidence consistent with the interpretation that the effect reflects active, encoding-specific memory reprocessing rather than a stable individual sleep characteristic. These findings extend the active systems consolidation framework to encompass visual pattern completion and complement existing evidence on sleep-dependent pattern separation (Hanert et al., 2017) and verbal associative completion (Lutz et al., 2024), suggesting distinct oscillatory mechanisms by which NREM sleep shapes hippocampal representations for flexible memory-based generalisation.

## Acknowledgements

This study has been supported by the German Research Foundation (DFG) FOR 5434, by the Else Kröner-Fresenius-Stiftung and by the Faculty of Medicine, University of Kiel, Germany.

## Author contributions

T.B., A.H., and J.B. designed the study. A.H. performed the analyses, A.H. and T.B. interpreted the data, and drafted the manuscript. Y.F. and P.V. contributed to data analysis and interpretation. J.H. collected the data. A.B. and A.P. contributed to the conceptual development of the study, provided supervision, and critically revised the manuscript. All authors contributed to the final version of the manuscript and approved its submission.

## Data Availability Statement

The datasets generated and analyzed during the current study are available from the corresponding author on reasonable request.

## References

Amer, T., & Davachi, L. (2023). Extra-hippocampal contributions to pattern separation. ELife, 12. 10.7554/eLife.82250

Cellini, N., Mercurio, M., Vanzetti, V., Bergamo, D., & Sarlo, M. (2020). Comparing the effect of daytime sleep and wakefulness on mnemonic discrimination. Physiology & Behavior, 224, 113078. 10.1016/j.physbeh.2020.113078

Clemens, Z., Mölle, M., Eross, L., Barsi, P., Halász, P., & Born, J [Jan] (2007). Temporal coupling of parahippocampal ripples, sleep spindles and slow oscillations in humans. Brain : A Journal of Neurology, 130(Pt 11), 2868–2878. 10.1093/brain/awm146

Deuker, L., Doeller, C. F., Fell, J., & Axmacher, N. (2014). Human neuroimaging studies on the hippocampal CA3 region - integrating evidence for pattern separation and completion. Frontiers in Cellular Neuroscience, 8, 64. 10.3389/fncel.2014.00064

Diekelmann, S., & Born, J [Jan] (2010). The memory function of sleep. Nature Reviews. Neuroscience, 11(2), 114–126. 10.1038/nrn2762

Doxey, C. R., Hodges, C. B., Bodily, T. A., Muncy, N. M., & Kirwan, C. B. (2018). The effects of sleep on the neural correlates of pattern separation. Hippocampus, 28(2), 108–120. 10.1002/hipo.22814

Ellenbogen, J. M., Hu, P. T., Payne, J. D., Titone, D., & Walker, M. P. (2007). Human relational memory requires time and sleep. Proceedings of the National Academy of Sciences of the United States of America, 104(18), 7723–7728. 10.1073/pnas.0700094104

Fernandez, L. M. J., & Lüthi, A. (2020). Sleep Spindles: Mechanisms and Functions. Physiological Reviews, 100(2), 805–868. 10.1152/physrev.00042.2018

Grande, X., Berron, D., Horner, A. J., Bisby, J. A., Düzel, E., & Burgess, N. (2019). Holistic Recollection via Pattern Completion Involves Hippocampal Subfield CA3. The Journal of Neuroscience, 39(41), 8100–8111. 10.1523/JNEUROSCI.0722-19.2019

Guzman, S. J., Schlögl, A., Frotscher, M., & Jonas, P. (2016). Synaptic mechanisms of pattern completion in the hippocampal CA3 network. *Science (New York*, N.Y.), 353(6304), 1117–1123. 10.1126/science.aaf1836

Hanert, A., Weber, F. D., Pedersen, A., Born, J [J.], & Bartsch, T. (2017). Sleep in humans stabilizes hippocampal pattern separation performance. Clinical Neurophysiology, 129(8), e91. 10.1016/j.clinph.2018.04.695

Heib, D. P. J., Hoedlmoser, K., Anderer, P., Zeitlhofer, J., Gruber, G., Klimesch, W., & Schabus, M. (2013). Slow oscillation amplitudes and up-state lengths relate to memory improvement. PloS One, 8(12), e82049. 10.1371/journal.pone.0082049

Horner, A. J., Bisby, J. A., Bush, D., Lin, W.-J., & Burgess, N. (2015). Evidence for holistic episodic recollection via hippocampal pattern completion. Nature Communications, 6, 7462. 10.1038/ncomms8462

Hunsaker, M. R., & Kesner, R. P. (2013). The operation of pattern separation and pattern completion processes associated with different attributes or domains of memory. Neuroscience and Biobehavioral Reviews, 37(1), 36–58. 10.1016/j.neubiorev.2012.09.014

Inostroza, M., & Born, J [Jan] (2013). Sleep for preserving and transforming episodic memory. Annual Review of Neuroscience, 36, 79–102. 10.1146/annurev-neuro-062012-170429

Joensen, B. H., Ashton, J. E., Berens, S. C., Gaskell, M. G., & Horner, A. J. (2024). An Enduring Role for Hippocampal Pattern Completion in Addition to an Emergent Nonhippocampal Contribution to Holistic Episodic Retrieval after a 24 h Delay. The Journal of Neuroscience, 44(18). 10.1523/JNEUROSCI.1740-23.2024

Klinzing, J. G., Niethard, N., & Born, J [Jan] (2019). Mechanisms of systems memory consolidation during sleep. Nature Neuroscience, 22(10), 1598–1610. 10.1038/s41593-019-0467-3

Kumaran, D., Hassabis, D., & McClelland, J. L. (2016). What Learning Systems do Intelligent Agents Need? Complementary Learning Systems Theory Updated. Trends in Cognitive Sciences, 20(7), 512–534. 10.1016/j.tics.2016.05.004

Kumaran, D., & McClelland, J. L. (2012). Generalization through the recurrent interaction of episodic memories: A model of the hippocampal system. Psychological Review, 119(3), 573–616. 10.1037/a0028681

Latchoumane, C.-F. V., Ngo, H.-V. V., Born, J [Jan], & Shin, H.-S. (2017). Thalamic Spindles Promote Memory Formation during Sleep through Triple Phase-Locking of Cortical, Thalamic, and Hippocampal Rhythms. Neuron, 95(2), 424-435.e6. 10.1016/j.neuron.2017.06.025

Leutgeb, J. K., Leutgeb, S., Moser, M.-B., & Moser, E. I. (2007). Pattern separation in the dentate gyrus and CA3 of the hippocampus. Science (New York, N.Y.), 315(5814), 961–966. 10.1126/science.1135801

Leutgeb, S., Leutgeb, J. K., Treves, A [Alessandro], Moser, M.-B., & Moser, E. I. (2004). Distinct ensemble codes in hippocampal areas CA3 and CA1. *Science (New York*, N.Y.), 305(5688), 1295– 1298. 10.1126/science.1100265

Lewis, P. A., & Durrant, S. J. (2011). Overlapping memory replay during sleep builds cognitive schemata. Trends in Cognitive Sciences, 15(8), 343–351. 10.1016/j.tics.2011.06.004

Liu, K. Y., Gould, R. L., Coulson, M. C., Ward, E. V., & Howard, R. J. (2016). Tests of pattern separation and pattern completion in humans-A systematic review. Hippocampus, 26(6), 705–717. 10.1002/hipo.22561

Lutz, N. D., Martínez-Albert, E., Friedrich, H., Born, J [Jan], & Besedovsky, L. (2024). Sleep shapes the associative structure underlying pattern completion in multielement event memory. Proceedings of the National Academy of Sciences of the United States of America, 121(9), e2314423121. 10.1073/pnas.2314423121

Marr, D. (1971). Simple memory: A theory for archicortex. *Philosophical Transactions of the Royal Society of London. Series B*, Biological Sciences, 262(841), 23–81. 10.1098/rstb.1971.0078

McClelland, J. L., McNaughton, B. L [Bruce L.], & O’Reilly, R. C. (1995). Why there are complementary learning systems in the hippocampus and neocortex: Insights from the successes and failures of connectionist models of learning and memory. Psychological Review, 102(3), 419–457. 10.1037/0033-295X.102.3.419

McNaughton, B. L [B. L.], & Morris, R. (1987). Hippocampal synaptic enhancement and information storage within a distributed memory system. Trends in Neurosciences, 10(10), 408–415. 10.1016/0166-2236(87)90011-7

Mednick, S., Nakayama, K., & Stickgold, R. (2003). Sleep-dependent learning: A nap is as good as a night. Nature Neuroscience, 6(7), 697–698. 10.1038/nn1078

Mölle, M., Yeshenko, O., Marshall, L., Sara, S. J., & Born, J [Jan] (2006). Hippocampal sharp wave-ripples linked to slow oscillations in rat slow-wave sleep. Journal of Neurophysiology, 96(1), 62– 70. 10.1152/jn.00014.2006

Muehlroth, B. E., Sander, M. C., Fandakova, Y., Grandy, T. H., Rasch, B., Shing, Y. L., & Werkle-Bergner, M. (2019). Precise Slow Oscillation-Spindle Coupling Promotes Memory Consolidation in Younger and Older Adults. Scientific Reports, 9(1), 1940. 10.1038/s41598-018-36557-z

Nakazawa, K., Quirk, M. C., Chitwood, R. A., Watanabe, M., Yeckel, M. F., Sun, L. D., Kato, A., Carr, C. A., Johnston, D., Wilson, M. A., & Tonegawa, S. (2002). Requirement for hippocampal CA3 NMDA receptors in associative memory recall. *Science (New York*, N.Y.), 297(5579), 211–218. 10.1126/science.1071795

Neunuebel, J. P., & Knierim, J. J. (2014). Ca3 retrieves coherent representations from degraded input: Direct evidence for CA3 pattern completion and dentate gyrus pattern separation. Neuron, 81(2), 416–427. 10.1016/j.neuron.2013.11.017

Ng, T., Noh, E., & Spencer, R. M. C. (2025). Bayesian meta-analysis reveals the mechanistic role of slow oscillation-spindle coupling in sleep-dependent memory consolidation. ELife, 13. 10.7554/eLife.101992

Niknazar, M., Krishnan, G. P., Bazhenov, M., & Mednick, S. C. (2015). Coupling of Thalamocortical Sleep Oscillations Are Important for Memory Consolidation in Humans. PloS One, 10(12), e0144720. 10.1371/journal.pone.0144720

Norman, K. A., & O’Reilly, R. C. (2003). Modeling hippocampal and neocortical contributions to recognition memory: A complementary-learning-systems approach. Psychological Review, 110(4), 611–646. 10.1037/0033-295X.110.4.611

Rasch, B., & Born, J [Jan] (2013). About sleep’s role in memory. Physiological Reviews, 93(2), 681–766. 10.1152/physrev.00032.2012

Rasch, B., Büchel, C., Gais, S., & Born, J [Jan] (2007). Odor cues during slow-wave sleep prompt declarative memory consolidation. *Science (New York*, N.Y.), 315(5817), 1426–1429. 10.1126/science.1138581

Rave, J., Hanert, A., Tabi, Y. A., Fiedler, Y., Philippen, S., Granert, O., Leypoldt, F., Born, J [J.], Burgalossi, A., Schoenfeld, R., Finke, C., & & Bartsch, T. (2026). Impaired consolidation of spatial memory during sleep in patients with LGI1-associated limbic encephalitis. Brain Communications(accepted for publication).

Reichardt, R., Király, A., Szőllősi, Á., Racsmány, M., & Simor, P. (2024). A daytime nap with REM sleep is linked to enhanced generalization of emotional stimuli. Journal of Sleep Research, 33(5), e14177. 10.1111/jsr.14177

Rolls, E. T [Edmund T.] (2013). The mechanisms for pattern completion and pattern separation in the hippocampus. Frontiers in Systems Neuroscience, 7, 74. 10.3389/fnsys.2013.00074

Rolls, E. T [Edmund T.] (2016). Pattern separation, completion, and categorisation in the hippocampus and neocortex. Neurobiology of Learning and Memory, 129, 4–28. 10.1016/j.nlm.2015.07.008

Staresina, B. P., Bergmann, T. O., Bonnefond, M., van der Meij, R., Jensen, O., Deuker, L., Elger, C. E., Axmacher, N., & Fell, J. (2015). Hierarchical nesting of slow oscillations, spindles and ripples in the human hippocampus during sleep. Nature Neuroscience, 18(11), 1679–1686. 10.1038/nn.4119

Stark, S. M., Yassa, M. A., Lacy, J. W., & Stark, C. E. L. (2013). A task to assess behavioral pattern separation (BPS) in humans: Data from healthy aging and mild cognitive impairment. Neuropsychologia, 51(12), 2442–2449. 10.1016/j.neuropsychologia.2012.12.014

Treves, A [A.], & Rolls, E. T [E. T.] (1994). Computational analysis of the role of the hippocampus in memory. Hippocampus, 4(3), 374–391. 10.1002/hipo.450040319

Vieweg, P., Riemer, M., Berron, D., & Wolbers, T. (2019). Memory Image Completion: Establishing a task to behaviorally assess pattern completion in humans. Hippocampus, 29(4), 340–351. 10.1002/hipo.23030.

Vieweg, P., Stangl, M., Howard, L. R., & Wolbers, T. (2015). Changes in pattern completion--a key mechanism to explain age-related recognition memory deficits? Cortex; a Journal Devoted to the Study of the Nervous System and Behavior, 64, 343–351. 10.1016/j.cortex.2014.12.007

Walker, M. P., & Stickgold, R. (2010). Overnight alchemy: Sleep-dependent memory evolution. Nature Reviews. Neuroscience, 11(3), 218; author reply 218. 10.1038/nrn2762-c1

Werchan, D. M., & Gómez, R. L. (2013). Generalizing memories over time: Sleep and reinforcement facilitate transitive inference. Neurobiology of Learning and Memory, 100, 70–76. 10.1016/j.nlm.2012.12.006

Witkowski, S., Noh, S., Lee, V., Grimaldi, D., Preston, A. R., & Paller, K. A. (2021). Does memory reactivation during sleep support generalization at the cost of memory specifics? Neurobiology of Learning and Memory, 182, 107442. 10.1016/j.nlm.2021.107442

Yassa, M. A., & Stark, C. E. L. (2011). Pattern separation in the hippocampus. Trends in Neurosciences, 34(10), 515–525. 10.1016/j.tins.2011.06.006

Zeithamova, D., & Bowman, C. R. (2020). Generalization and the hippocampus: More than one story? Neurobiology of Learning and Memory, 175, 107317. 10.1016/j.nlm.2020.107317

