## Supplemental Table 1 for "Post-encoding slow-wave amplitude during a daytime nap predicts pattern completion from sparse visual cues"

**Table S1.** Descriptive statistics for pattern completion, pattern separation, and the bias measure before and after the sleep and wake retention intervals across completeness levels.

| | Pattern Completion (Learned Stimuli) | | | | Pattern Separation (New Stimuli) | | | | Bias measure | | | |
|---|---|---|---|---|---|---|---|---|---|---|---|---|
| Completeness (%) | Pre Sleep | Post Sleep | Pre Wake | Post Wake | Pre Sleep | Post Sleep | Pre Wake | Post Wake | Pre Sleep | Post Sleep | Pre Wake | Post Wake |
| 100 | 99.6 ± 1.6 | 99.3 ± 2.0 | 99.4 ± 1.9 | 99.4 ± 2.6 | 99.8 ± 1.2 | 100.0 ± 0 | 99.8 ± 1.2 | 99.6 ± 1.6 | -0.2 ± 2.1 | -0.7 ± 2.0 | -0.4 ± 2.3 | -0.2 ± 3.1 |
| 35 | 94.9 ± 6.6 | 99.1 ± 2.8 | 93.8 ± 8.5 | 97.8 ± 6.3 | 96.4 ± 4.0 | 100.0 ± 0 | 97.8 ± 3.8 | 100.0 ± 0 | -1.6 ± 6.7 | -0.9 ± 2.8 | -4.1 ± 8.7 | -2.2 ± 6.3 |
| 21 | 93.3 ± 8.1 | 97.3 ± 5.2 | 92.2 ± 9.1 | 97.4 ± 6.8 | 94.9 ± 5.9 | 100.0 ± 0 | 96.8 ± 4.3 | 99.6 ± 1.6 | -1.6 ± 10.0 | -2.7 ± 5.2 | -4.5 ± 10.9 | -2.2 ± 6.3 |
| 12 | 85.9 ± 11.9 | 94.9 ± 6.6 | 86.6 ± 11.8 | 95.9 ± 6.7 | 92.9 ± 8.6 | 99.3 ± 2.6 | 95.0 ± 5.6 | 98.7 ± 4.4 | -6.9 ± 13.0 | -4.5 ± 7.0 | -8.4 ± 12.1 | -2.8 ± 5.4 |
| 5 | 73.2 ± 19.1 | 88.6 ± 10.8 | 75.9 ± 16.6 | 82.3 ± 13.7 | 88.6 ± 14.8 | 96.4 ± 6.2 | 89.0 ± 15.5 | 94.8 ± 8.8 | -15.4 ± 23.4 | -7.8 ± 11.0 | -13.1 ± 19.2 | -12.5 ± 14.9 |

*Note.* Values are presented as mean ± SD. Completeness refers to the proportion of the original scene information available in the degraded retrieval cue (100%, 35%, 21%, 12%, or 5%). Pattern completion refers to the percentage of correctly identified learned stimuli, whereas pattern separation refers to the percentage of correctly identified new stimuli. The bias measure was calculated as pattern completion minus pattern separation. Pre (R1) and post (R2) denote performance before and after the 90-min sleep or wake retention interval.
